# Intestine-to-neuronal signaling alters risk-taking behaviors in food-deprived *Caenorhabditis elegans*

**DOI:** 10.1101/156109

**Authors:** Molly A. Matty, Hiu E. Lau, Anupama Singh, Jessica A. Haley, Ahana Chakraborty, Karina Kono, Kirthi C. Reddy, Malene Hansen, Sreekanth H. Chalasani

**Affiliations:** Molecular Neurobiology Laboratory, The Salk Institute for Biological Studies, La Jolla, CA 92037, USA; Division of Biological Sciences, University of California, San Diego, La Jolla, CA 92093, USA; Development, Aging and Regeneration Program, Sanford Burnham Prebys Medical Discovery Institute, La Jolla, CA 92037, USA; Neurosciences Graduate Program, University of California, San Diego, La Jolla, CA 92093, USA

**Keywords:** Behavioral plasticity, sensory integration, food-deprivation, MML-1, HLH-30, insulin-like peptides, DAF-2 receptors, ASI neurons, gut-brain axis.

## Abstract

Animals integrate changes in external and internal environments to generate behavior. While neural circuits detecting external cues have been mapped, less is known about how internal states like hunger are integrated into behavioral outputs. We use the nematode *C. elegans* to decode how changes in internal nutritional status affects chemosensory behaviors. We show that acute food deprivation leads to a reversible decline in repellent, but not attractant, sensitivity. This behavioral change requires two conserved transcription factors MML-1 (Mondo A) and HLH-30 (TFEB), both of which translocate from the intestinal nuclei to the cytoplasm upon food deprivation. Next, we identify insulin-like peptides INS-23 and INS-31 as candidate ligands relaying food-status signals from the intestine to other tissues. Furthermore, we show that ASI chemosensory neurons use the DAF-2 insulin receptor, PI-3 Kinase, and the mTOR complex to integrate these intestine-released peptides. Together, our study shows how internal food status signals are integrated by transcription factors and intestine-neuron signaling to generate flexible behaviors.

**Author Summary:** We have all experienced behavioral changes when we are hungry - the pang in our stomach can cause us to behave erratically. In particular, hungry animals, including humans, are known to pursue behaviors that involve higher risk compared to when they are well-fed. Here we explore the molecular details of this behavior in the invertebrate animal model C. elegans. This behavior, termed sensory integration, shows that C. elegans display reduced copper sensitivity when hungry. Copper is toxic and repellant to C. elegans; reduced avoidance indicates that these animals use riskier food search behaviors when they are hungry. Luckily, like us, this behavioral change is reversible upon re-feeding. This hunger-induced behavioral change is not due to increased attraction to food or depletion of fat stores, but rather insulin signaling between the intestine and specific neurons. We use genetic tools, microscopy, and behavioral tests to determine that this risky behavior involves sensation of “lack of food” in the intestine, release of signaling molecules, and engagement with sensory neurons. Our work highlights new and potentially evolutionarily conserved ways in which intestinal cells and neurons communicate leading to largescale behavioral change, providing further support for the importance of the gut-brain-axis.

## Introduction

Animals evaluate their environment, integrating prior experiences and internal state information to optimize their behaviors in order to maximize rewards and avoid threats [1]. Moreover, changes in internal states play a critical role in adjusting the animal’s responses to external stimuli [2, 3]. One critical internal state is hunger, which has a profound effect on animal survival and elicits dramatic changes in food-seeking behaviors [2, 4]. Multiple species, including humans, have been shown to alter their chemosensory behavior during periods of starvation [5–10]. Despite this, less is known about how the nervous system integrates information about hunger status.

The nematode *Caenorhabditis elegans*, with just 302 neurons [11], and 20 cells in its intestine [12], provides a unique opportunity for a high-resolution analysis of how the nervous system integrates internal signals. Previous studies have shown that *C. elegans*, similar to mammals, exhibits a number of behavioral, physiological, and metabolic changes in response to altered nutritional status. *C. elegans* hermaphrodites retain eggs [13], are unlikely to mate with males [14], initiate altered foraging behaviors [15–17], and change their responses to environmental CO_2_ [18], salt [19], and pheromones [20] upon food deprivation. Moreover, many molecules that signal hunger are conserved between *C. elegans* and vertebrates. For example, neuropeptide Y (NPY) signaling influences feeding behaviors in nematodes and mammals [21–23]. Similar effects are also seen with insulin and dopamine signaling, which seem to act via modifying chemosensory activity and behavior in nematodes [24, 25]; and on mammalian neural circuits [26–28] to modify feeding behavior. While neuronal pathways responding to food-deprivation on the multiple-minute timescales have been mapped [17, 29], those integrating these signals on the multiple hour timescales are poorly understood.

Here we use *C. elegans* to dissect the machinery required to integrate internal food signals and modify behaviors. We combined food deprivation over multiple hours with a behavioral assay that quantifies the animal’s ability to integrate both toxic and food-related signals, mimicking a simplified ecologically relevant scenario. In this sensory integration assay, animals cross a toxic copper barrier (repellent) and chemotax towards a point source of a volatile food-associated odor, diacetyl (attractant) [30]. We show that animals’ food-deprived for multiple hours have reduced sensitivity to the repellent and cross the copper barrier more readily than well-fed animals. Next, we show that two transcription factors translocate from the intestinal nuclei to the cytoplasm upon multiple hours of food deprivation. We confirm a role for these transcription factors and identify the downstream peptides released by the intestine to relay “the lack of food” signal to other tissues. Finally, we show that ASI chemosensory neurons integrate these intestine-released peptides. This allows animals to reduce their avoidance to repellents and undertake a higher risk strategy in their search for food.

## Results

### Acute food deprivation specifically alters repellent-driven behaviors

Animals simultaneously integrate both attractant and repellent signals from their environment to generate appropriate behavioral readouts. To mimic these interactions, animals are exposed to a copper repellent barrier (CuSO_4_) and a gradient of a volatile attractant, diacetyl [30]. The proportion of animals that cross the copper barrier are counted and expressed as a chemotactic index (**Figure 1A**). We analyzed the behavior of well-fed, wild-type animals and found that ∼30% crossed the copper barrier and locomote towards the spot of diacetyl (black bars, **Figure 1B and Movie S1**). In contrast, food-deprived animals were more likely to cross the copper barrier (blue bars, **Figure 1B and Movie S2**). We also found that animals needed to be food deprived for at least 1 hr before they significantly altered their behavior with a maximal effect at 3 hrs (**Figure 1B**). Next, we tested whether the food-deprivation effect was reversible. We food-deprived animals for 3 hrs and then returned them to food for different durations and analyzed animal behavior after the food experience. We found that 3-hr food-deprived animals that had been returned to food for at least 3 hrs reverted to the “well-fed” state (**Figure 1B**). Taken together, these results indicate that food deprivation reversibly modifies sensory integration behavior.

**Figure 1:**
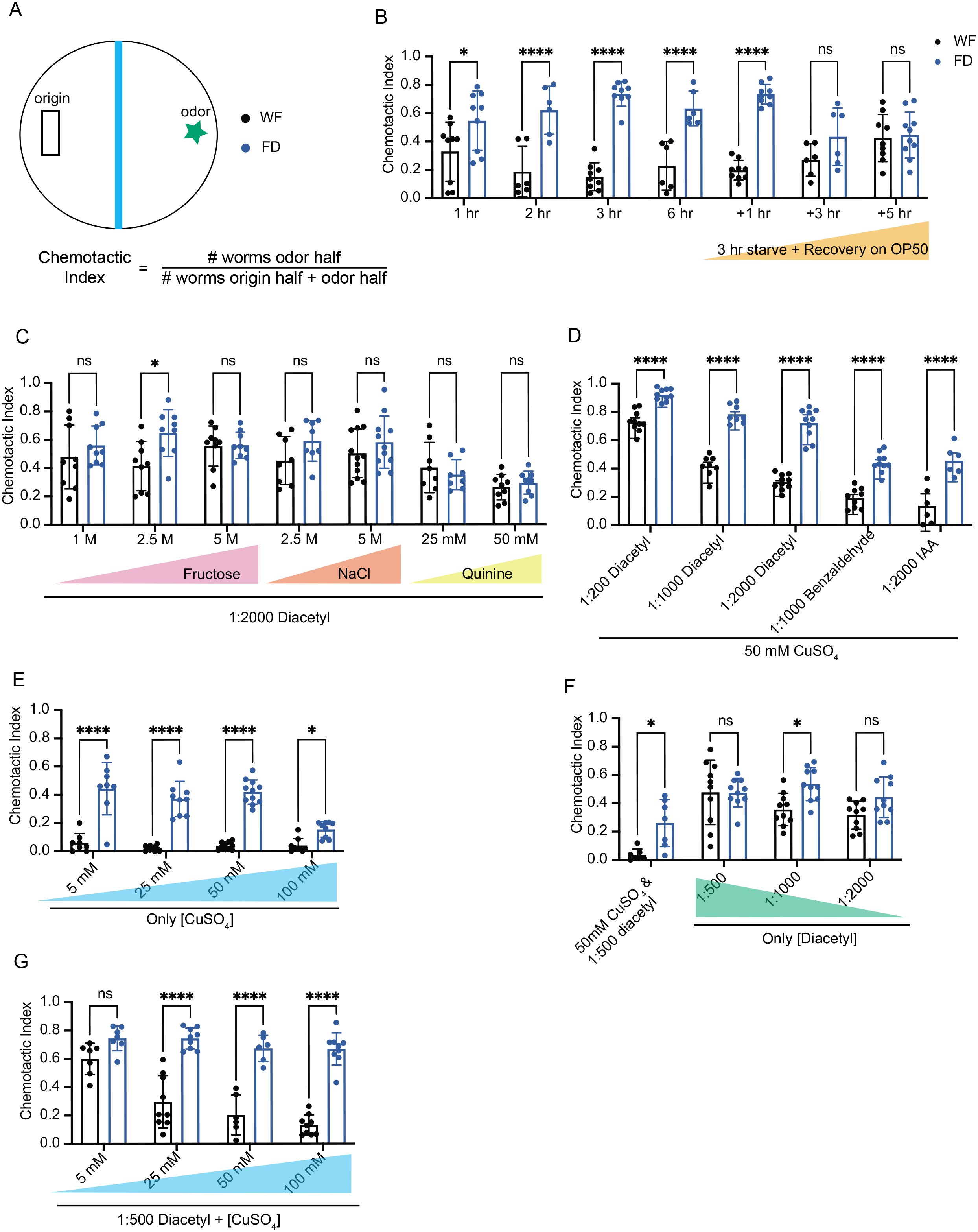
Starvation reduces copper avoidance. A) Schematic of the sensory integration assay. ∼100-200 day 1 adult animals (n) are placed in the black rectangle. Blue barrier represents copper barrier (or other repellant) and star represents diacetyl or other attractant. Chemotactic Index is the number of animals that have crossed the barrier (odor side) divided by the total number of animals on the plate (odor + origin sides). Experiments with well-fed (WF) animals will appear with black dots and those with food-deprived (FD) animals will be indicated with blue dots. Unless otherwise noted, FD is 3 hours with no food. Each dot represents a single plate (N) of animals (n). B) Animals are deprived of food for increasing periods of time. Animals that have been starved for 3 hours are allowed to recover for 1, 3, or 5 hours on OP50. Well-fed matched partners are kept on OP50 plates for the entire length of the experiment. Animals are exposed to 50 mM CuSO_4_ repellant and 1:500 (0.2%) diacetyl attractant. N≥6. C) Animals are exposed to increasing concentrations of other repellants (Fructose, NaCl, Quinine) with the attractant 0.05% diacetyl (1:2000) in each condition N≥7. D) Animals are exposed to decreasing concentrations of diacetyl (0.2%, 0.1% and 0.05%, or 1:200, 1:1000, and 1:2000, respectively) and other volatile attractants 0.1% Benzaldehyde (BZ) and 0.05% Isoamyl Alcohol (IAA). 50 mM CuSO_4_ is the repellant in each condition N≥6. E) Animals are exposed to CuSO_4_ in increasing concentrations (5 mM, 25 mM, 50 mM, 100 mM) without any attractant N≥8. F) Animals are exposed to diacetyl alone in decreasing concentrations (0.2%, 0.1%, 0.05%). Full assay (0.2% diacetyl and 50 mM CuSO_4_) is included as a control N≥7. G) Animals are exposed to 1:500 diacetyl and increasing concentrations of CuSO_4_ (5 mM, 25 mM, 50 mM, 100 mM) N≥6. All graphs are analyzed using a two-way ANOVA, determined to have significant differences across well-fed and food-deprived conditions. WF/FD comparisons were then performed as pairwise comparisons within each genotype or treatment as t-tests with Bonferroni corrections for multiple comparisons. * p<0.5, ** p<0.01, *** p<0.001, **** p<0.0001, ns p>0.05. Error bars are S.D.

We then tested whether this food deprivation-evoked change in sensory integration behavior was specific to the copper repellent and diacetyl attractant used in the assay. We observed that food-deprived animals did not cross the repellent barrier when diacetyl was paired with other repellents like fructose (except one intermediate concentration), sodium chloride, or quinine (**Figure 1C**). In contrast when copper was paired other attractants like benzaldehyde and isoamyl alcohol, food deprived animals continued to cross the copper barrier more readily than well-fed animals (**Figure 1D**). Next, we tested responses of these animals to varying concentrations of copper or diacetyl alone. We found that food-deprived animals crossed the copper barrier more readily than well-fed animals, suggesting that their responsiveness to copper is reduced even in the absence of an attractant (**Figure 1E**). In contrast, food-deprived animals did not discernably alter their attraction to diacetyl in the absence of the copper repellant (**Figure 1F**). Given the small number of well-fed animals that cross the copper barrier alone (**Figure 1E**), we continued to pair copper with the diacetyl attractant for further analysis. We also tested whether altering the concentrations of the copper barrier has an effect on food-deprived animals and confirmed these concentrations using a copper indicator (**Supplementary Figure 1**). We found that food-deprived animals showed significant increase in their ability to cross the repellent barrier above a threshold of 5 mM copper concentration (**Figure 1G**). To gain further confirmation of this copper-specific change, we tested food-deprived animals in a single animal copper drop assay (**Supplementary Figure 2A**). In this assay, the response of a single animal to a drop of 1.5 mM CuSO_4_ solution placed in its path was monitored. Most repellents can be tested in this assay with animals generating a robust avoidance response [31]. We found that food-deprived animals showed a significant deficit in their copper avoidance response (**Supplementary Figure 2B**). Collectively, these data show that food-deprived animals display reduced sensitivity to copper, which we dissected further using genetic methods and tracking software.

### Dynamics of risky search strategies in food-deprived animals

To analyze how food deprivation modifies animal behavior, we recorded and tracked populations of animals over 45 minutes in the sensory integration assay. Individual animal trajectories were identified and used for analysis (see Methods and **Supplementary Figure 3**). We found that fewer well-fed animals cross the repellent copper barrier (**Figure 2A**) as compared to food-deprived animals (**Figure 2B**) during the entire 45 min assay (example tracks for all groups in **Supplementary Figure 3 E-H**). To quantify this difference, we plotted the fraction of tracks that crossed the copper barrier as a function of time (**Figure 2C**). We found that food-deprived, wild-type animals were more likely to cross the barrier at all time points (15, 30, and 45 mins) compared to well-fed animals. Thus, the differences between well-fed and food-deprived animals were not limited to specific time windows in the assay. To further assess the increased likelihood of food-deprived animals crossing the repellent barrier, we compared the probability of animal tracks being located at given distances from the barrier (**Figure 2D,** methods described in **Supplementary Figure 3A-C**)). We found that food-deprived worms are nearly twice as likely to reside within +/-0.5 cm from the copper barrier while well-fed animals are more likely to be found 2.1 cm from the barrier, not far from where the animals were placed on the assay plate (**Figure 2D**, statistics summarized in **Supplementary Table 2**). These data suggest that well-fed animals reorient upon detection of the copper thereby increasing the likelihood of animals being located in regions well before the barrier. In contrast, food-deprived animals cross the barrier more frequently. To further dissect these behavioral differences, we quantified the mean velocity of worm tracks (**Figure 2E**). We find that well-fed animals move more slowly when physically closer to the copper barrier. Food-deprived worms are significantly slower at distances far from the copper barrier (2-3.5 cm), but then accelerate to speeds matching well-fed behavior as they approach the barrier before slowing down as they reach the barrier consistent with well-fed animals (**Figure 2E, Supplementary Table 2**).

**Figure 2:**
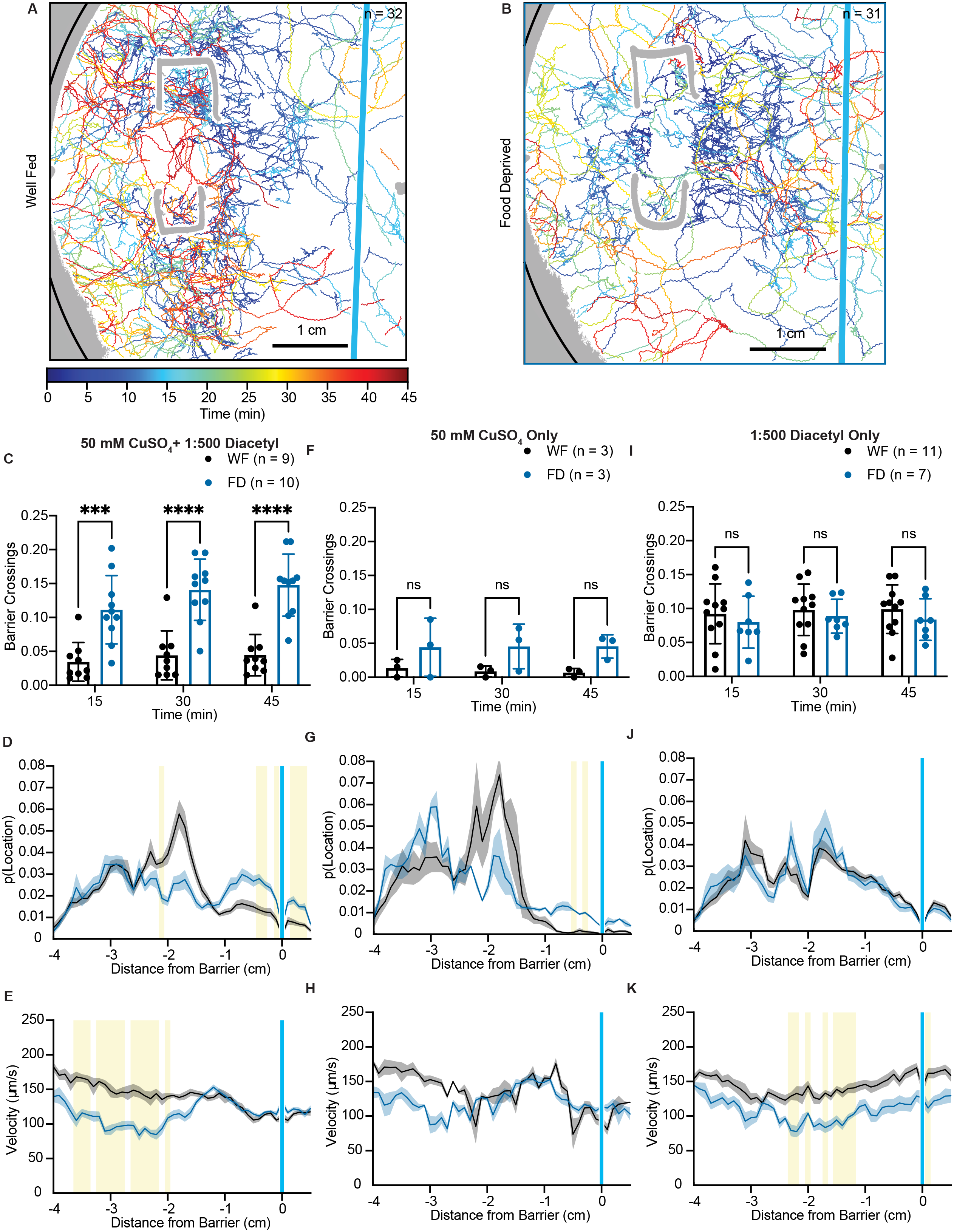
Riskier search strategies in starved worms. (A) Worm tracks (n = 32) are plotted for a representative sensory integration assay of well-fed worms behaving in the presence of 50mM CuSO_4_ (blue stripe) and 1 μL 0.2% diacetyl (1:500) (location not shown). Regions of the plate that were not able to be tracked are in gray with the edge of the plate indicated in black. Tracks are plotted and color coded for time (0 – 45 minutes). (B) Worm tracks (n = 31) are plotted for a representative sensory integration assay of 3 hour food-deprived worms. Conditions and plotting the same as in A. (C, F, I) The mean cumulative sum of worm tracks that cross the barrier as a fraction of the total worm tracks is plotted at three time points (15, 30, and 45 minutes). Well-fed (WF) animals appear with black dots and food-deprived (FD) animals are indicated with blue dots. Each dot represents a single plate of animals. C) 50 mM CuSO_4_ and 0.2% diacetyl F) 50 mM CuSO_4_, no diacetyl I) No copper, 0.2% (1:500) diacetyl. Graphs are analyzed using a two-way ANOVA to determine significant differences across well-fed and food-deprived conditions. WF/FD comparisons were then performed as pairwise comparisons within each time period as t-tests with Bonferroni corrections for multiple comparisons. * p<0.5, ** p<0.01, *** p<0.001, **** p<0.0001, ns p>0.05. (D, G, J) The probability of an animal being located at 1 mm binned distances from the barrier is plotted for well-fed (black) and food-deprived animals (blue). The dark line represents the mean probability of residence with the shaded areas representing the standard error of the mean. D) 50 mM CuSO_4_ and 0.2% diacetyl G) 50 mM CuSO_4_, no diacetyl J) No copper, 0.2% diacetyl. For each graph, multiple unpaired t-tests with Welch’s correction were performed with correction for multiple comparisons with Holm-Šídák post-hoc test. Corrected p values <0.05 are indicated by yellow shading. A comprehensive list of the statistics can be found in Supplementary Table 2. (E, H, K) The mean velocity of worms as a function of distance from the barrier is plotted for well-fed (black) and food-deprived animals (blue). Conditions, plotting, and statistics are the same as in D, G, and J.

When this assay is run in the absence of the attractant diacetyl, there is no significant difference in the probability of crossing the copper barrier between well-fed and food-deprived worms (**Figure 2F**), but this may be related to the low numbers of experiments we analyzed in this group. However, this is consistent with an increased chemotactic index of food-deprived worms across the copper barrier in the absence of an attractant at all time points (**Supplementary Figure 3D**). The probability of food-deprived animals locomoting close to the copper barrier is higher for 0.3 and 0.5 cm before the barrier **(Figure 2G, Supplementary Table 2**) while the velocity of these worms as a function of distance to the barrier is distributed similarly to worms assayed with copper and diacetyl (**Figure 2H, Supplementary Table 2).** These data suggest that the increased likelihood of food-deprived animals to cross the copper barrier cannot be explained by a deficiency in copper sensation alone. Further, these data suggest that food-deprived animals display similar locomotion dynamics to well-fed animals in response to copper in the absence of diacetyl.

In the absence of copper, food-deprived and well-fed animals behave similarly with no significant difference in the fraction of “barrier” (no copper) crossings toward the diacetyl or their localization on the assay plate (**Figure 2I, 2J, Supplementary Table 2**). Further, food-deprived worms display a decreased velocity on average (**Figure 2K, Supplementary Table 2**), consistent with previous studies [32]. Collectively, these data suggest that increase in food-deprived animals crossing the copper barrier is not due increased mobility, but may rather be a result of these animals pursuing navigation that is unfavorable (copper is toxic to *C. elegans* [33]).

### Lack of food and not changes in fat drives the food-deprivation induced behavioral change

Given that the change in sensory integration behavior requires multiple hours of food-deprivation, we hypothesized that metabolic signals like changes in fat content might play a crucial role. Additionally, previous studies have shown that prolonged starvation can deplete fat stores in *C. elegans*, which in turn can affect behavior [34, 35]. We tested whether 3 hrs of food deprivation alters the fat content of animals. Oil-Red O (ORO), a fat-soluble dye that stains triglycerides and lipoproteins has been used to label and quantify fat stores in *C. elegans* (**Figure 3A-B**) [36]. We used this dye and found that 3 hrs of food-deprivation did not alter the ORO signal or the area of the animal labeled by this stain (**Figure 3C-D**). In contrast, we observed a significant change in the both the intensity of the signal and area of animal stained in 6-hr food-deprived animals, consistent with previous studies [37]. These data suggest that changes in sensory integration behavior, which occurs after 3 hrs of food-deprivation is likely to be independent of fat metabolism.

**Figure 3:**
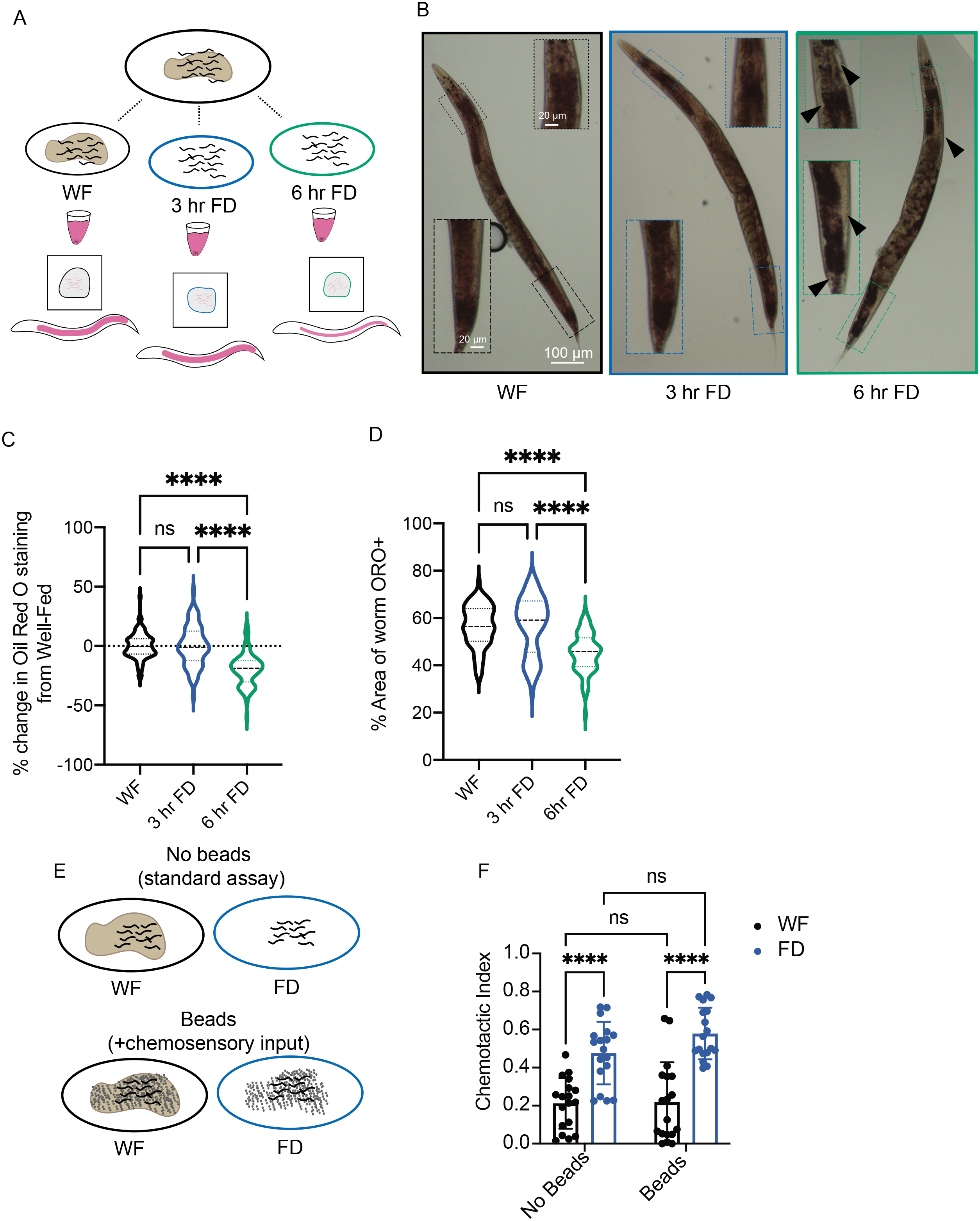
Lack of food, not fat or physical interactions, drive behavioral changes. (A) Schematic of Oil Red O experiments. Animals are raised together to day 1 of adulthood and separated into three groups: well-fed (on food), 3 hour food-deprived, and 6 hour food-deprived. Animals are stained using Oil Red O and then imaged using a color camera. (B) Representative images of well-fed (WF, black), 3 hour food-deprived (3hr FD, blue), and 6 hour food-deprived (6hr FD, green). Inset images are shown, highlighting the regions where there is the most difference in staining. Black arrows highlight regions of no Oil Red O stain in 6hr FD. (C) Graph showing the percent change in Oil Red O staining when compared to the average of the area of Oil Red O signal above a threshold value in the well-fed group within each independent experiment. N=3, n>20 within each experimental treatment group. (D) Graph showing the percent of the animals’ area that contains Oil Red O signal above threshold N=3, n>20 within each experimental treatment group. Same data as in C, shown as non-normalized values. (E) A schematic representing the experiment in F, in which populations of animals are either well-fed or food-deprived in the presence or absence of Sephadex beads before performing the sensory integration assay. (F) Prior to the sensory integration assay, animals are exposed to either standard OP50 (“no beads WF”) or empty plates (“no beads FD”), or Sephadex gel beads as chemosensory input. Alternatively, animals were exposed to beads and no food (“beads FD”) or OP50 with Sephadex beads on top (“beads WF”) for 3 hours. Animals were then exposed to standard Sensory Integration Assay set-up with 50 mM CuSO_4_ and 1 μL of 0.2% diacetyl. N≥18. C and D were analyzed using Welch’s ANOVA test with Dunnett’s multiple comparisons test. * p<0.5, ** p<0.01, *** p<0.001, **** p<0.0001, ns p>0.05. F was analyzed using a full model two-way ANOVA, determined to have significant differences across well-fed and food-deprived conditions but no difference between “bead”/“no bead” groups. Those comparisons are shown to indicate no difference between “beads” and “no beads”. Pairwise comparisons within each treatment were performed as t-tests with Tukey’s multiple comparisons test. Error bars are S.D.

Next, we sought to identify the relevant aspects of the bacterial experience contribute to the food deprivation-triggered behavioral change. *C. elegans* has been shown to evaluate multiple aspects of the food experience, including changes in food distribution, oxygen and carbon dioxide concentrations, small molecule metabolites and others [38–40]. To uncouple the tactile and chemosensory input of the bacteria from the nutritional value of ingesting bacteria, we analyzed the effect of using Sephadex gel beads on animal behavior. Animals exposed to gel beads experience the tactile input, but are not exposed to the nutritional value of food (**Figure 3E**) [15]. Notably, we found that animals exposed o Sephadex beads in the absence of *E. coli* OP50 for 3 hrs behaved the same as food-deprived animals in the sensory integration assay (**Figure 3F**). Together, these results show that the lack of food in the *C. elegans* intestine, but not the absence of chemosensory cues, reduces the animal’s sensitivity to copper.

### Transcription factors mediate food deprivation-induced behavioral change

Our studies indicated that the lack of food inside the animal was responsible for the transient reduction in copper sensitivity. To gain insights into the underlying molecular machinery, we investigated the role of nutritional-responsive transcription factors in the sensory integration assay. In mammalian cells, glucose is rapidly converted to glucose-6-phostphate, whose levels are sensed by a two basic-helix-loop-helix-leucine zipper transcription factors, MondoA and ChREBP (Carbohydrate Response Element Binding Protein). In well-fed conditions, MondoA binds the excess glucose-6-phosphate and Mlx (Max-like protein X) and translocates to the nucleus where it activates transcription of glucose-responsive genes. In the absence of glucose, MondoA remains in the cytoplasm [41, 42] (**Figure 4A**). *C. elegans* orthologs of MondoA and Mlx have been identified as MML-1 and MXL-2, respectively [43]. Furthermore, MML-1/MondoA has also been shown to translocate into the intestinal nuclei under well-fed conditions (**Figure 4A**) [44]. We predicted that *mml-1* mutants would be unable to sense the lack of food and thereby unable to reduce copper sensitivity after food deprivation. Consistently, we found that *mml-1*, but not *mxl-2* mutants were defective in their integration responses after food deprivation (**Figure 4B**). We then tested whether food deprivation alters the sub-cellular localization of the MML-1 protein. We monitored the GFP fluorescence under well-fed and food-deprived conditions in a *mml-1* knockout transgenic animal expressing GFP fused to the full-length coding sequence of MML-1/MondoA under well-fed and food-deprived conditions. We found that 3 hrs of food-deprivation resulted in an increased translocalization of MML-1/MondoA from the nucleus to the cytoplasm of the intestinal cells (**Figure 4C, 4D**). We suggest that this cytosolic MML-1/MondoA reduces copper sensitivity by modifying signaling between tissues.

**Figure 4:**
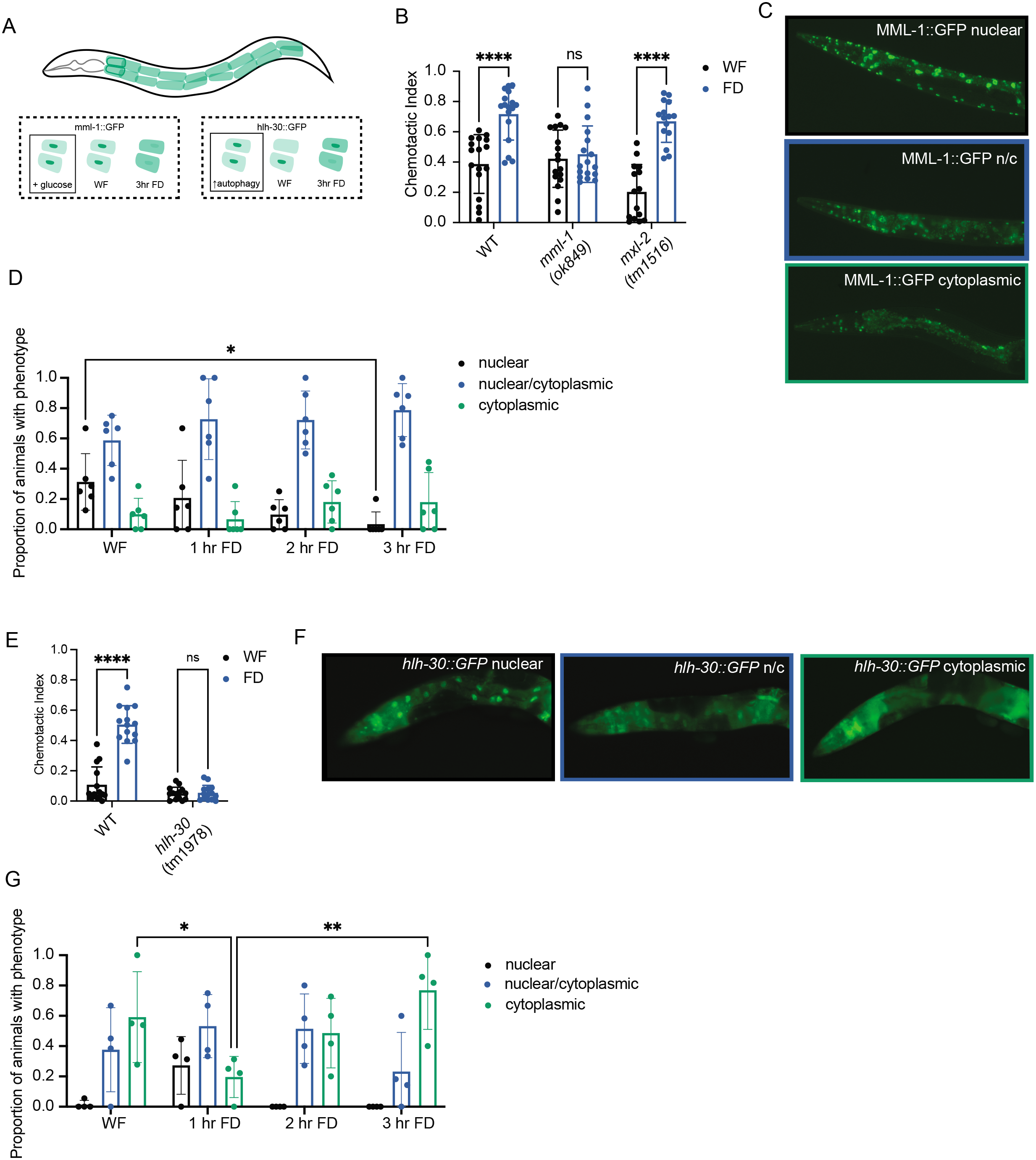
*mml-1* and *hlh-30* are required for sensory integration shift upon food deprivation, correlated with shifts in intestinal localization. (A) Schematic showing the 20 intestinal cells in a day 1 adult *C. elegans*. Our findings for MML-1:GFP and HLH-30::GFP are shown in the dotted box, while previously published paradigms are within the solid line box. Addition of glucose has been shown to induce nuclear localization of MondoA. Autophagy has been shown to increase nuclear localization of HLH-30. (B) Standard sensory integration assay with *mml-1(ok849)* and *mxl-2(tm1516* and wildtype controls. N=20. (C) Representative images of MML-1::GFP localization in day 1 adult animals (data quantified in D). All images were collected with the same exposure time and laser power. (D) Intestinal MML-1::GFP expression in animals during static timepoints food deprivation. Only intestinal expression was characterized as “nuclear”, “nuclear/cytoplasmic”, or “cytoplasmic”. Each dot represents the proportion of animals within an experiment with the phenotype. N=6, n=296. (E) Standard sensory integration assay with *hlh-30(tm1978)* mutant animals and wildtype controls. N=9. (F) Representative images of HLH-30::GFP localization in day 1 adult animals (data quantified in G). All images were collected with the same exposure time and laser power. (G) Intestinal HLH-30::GFP expression in animals during static timepoints of food deprivation. Only intestinal expression was characterized as “nuclear”, “nuclear/cytoplasmic”, or “cytoplasmic”. Each dot represents the proportion of animals within an experiment with the phenotype. N=3, n=149. B and E were analyzed using two-way ANOVA, determined to have significant differences across well-fed and food-deprived conditions. WF/FD comparisons were then performed as pairwise comparisons within each genotype or treatment as t-tests with Bonferroni’s multiple comparisons test. D and G were analyzed using Two-Way ANOVA, determined to have significant differences across localization and an interaction between time of food deprivation and localization. Within each localization group, pairwise comparisons were performed across each time point and tested for significance using Tukey’s multiple comparisons test. * p<0.5, ** p<0.01, *** p<0.001, **** p<0.0001, ns p>0.05. Error bars are S.D.

Previous studies have shown that MML-1 regulates the activity and nuclear localization of a second bHLH transcription factor HLH-30 (*C. elegans* TFEB, **Figure 4A**) [45]. In multiple animal models, HLH-30 functions as a key regulator of longevity pathways by promoting autophagy and lysosome biogenesis [46–49]. We tested whether HLH-30/TFEB was also required for food deprivation-evoked change in sensory integration. We found that, unlike wild-type animals, *hlh-30* null mutants did not show a change in their behavior after food-deprivation in the sensory integration assay (**Figure 4E**). We then tested whether the subcellular localization of HLH-30/TFEB was also affected by food deprivation. We observed an initial decrease in cytosolic GFP fluorescence at 1 hr of food-deprivation in HLH-30::GFP transgenic animals (Lapierre et al., 2011). Subsequently, at 3 hr of food-deprivation we found a robust increase in cytosolic HLH-30::GFP fluorescence (**Figure 4F, 4G**). Collectively, these data show that both MML-1 and HLH-30 accumulate in the intestinal cytoplasm upon 3 hr of food-deprivation and are required for the consequent behavioral change in sensory integration.

### Intestine-to-neuron signaling involves insulin signaling

Previous studies have shown that the *C. elegans* intestine is a major site for the transcriptional regulation of insulin-like peptide genes in response to starvation [50]. In addition, HLH-30/TFEB has been shown to act upstream of the insulin-signaling pathway in regulating the expression of neuronal chemoreceptor genes [51]. The *C. elegans* genome encodes about 40 insulin-like peptides [52] and all of these ligands are thought to bind and signal via a single tyrosine kinase DAF-2 receptor [53]. We hypothesized that insulin-like peptides might also act downstream of HLH-30/TFEB in relaying food status signals from the intestine to other tissues. Consistent with our hypothesis, multiple insulin-like peptides including INS-3, INS-4, INS-6, INS-10, INS-17, INS-18, INS-23, and INS-31 contain HLH-30/TFEB binding sites in their promoters [51]. In addition, INS-7, INS-8 and INS-37 have been shown to affect the subcellular localization of HLH-30/TFEB in the *C. elegans* intestine after mating (**Figure 5A**). [54]. We tested mutants in these insulin-like peptide genes for their ability to alter their sensory integration behavior after food deprivation. We similarly tested animals with a semi-dominant mutation in *daf-28*(*sa191*), since this allele has been shown to the prevent other insulin-like peptides from binding the common DAF-2 insulin receptor [55–57]. We found that multiple insulin-like peptides including INS-23 (*tm1875*), INS-31 (*tm3543*) and DAF-28 (*sa191*) were unable to respond to food deprivation. Specifically, these mutant animals did not display an increased ability to cross the repellent copper barrier when food deprived (**Figure 5B**), implying that these might be candidate signals relaying food status signals. In contrast, *ins-7* (*tm2001*), *ins-8* (*tm4144*), and *ins-37* (*tm6061*) mutants were similar to wild-type animals in their ability to cross the copper barrier in both well-fed and food-deprived conditions (**Figure 5C**). Taken together, these data suggest that the intestine might release INS-23, INS-31 and DAF-28 (or other insulin-like peptides whose binding to the DAF-2 insulin receptor is blocked by DAF-28) to relay hunger information to other tissues.

**Figure 5:**
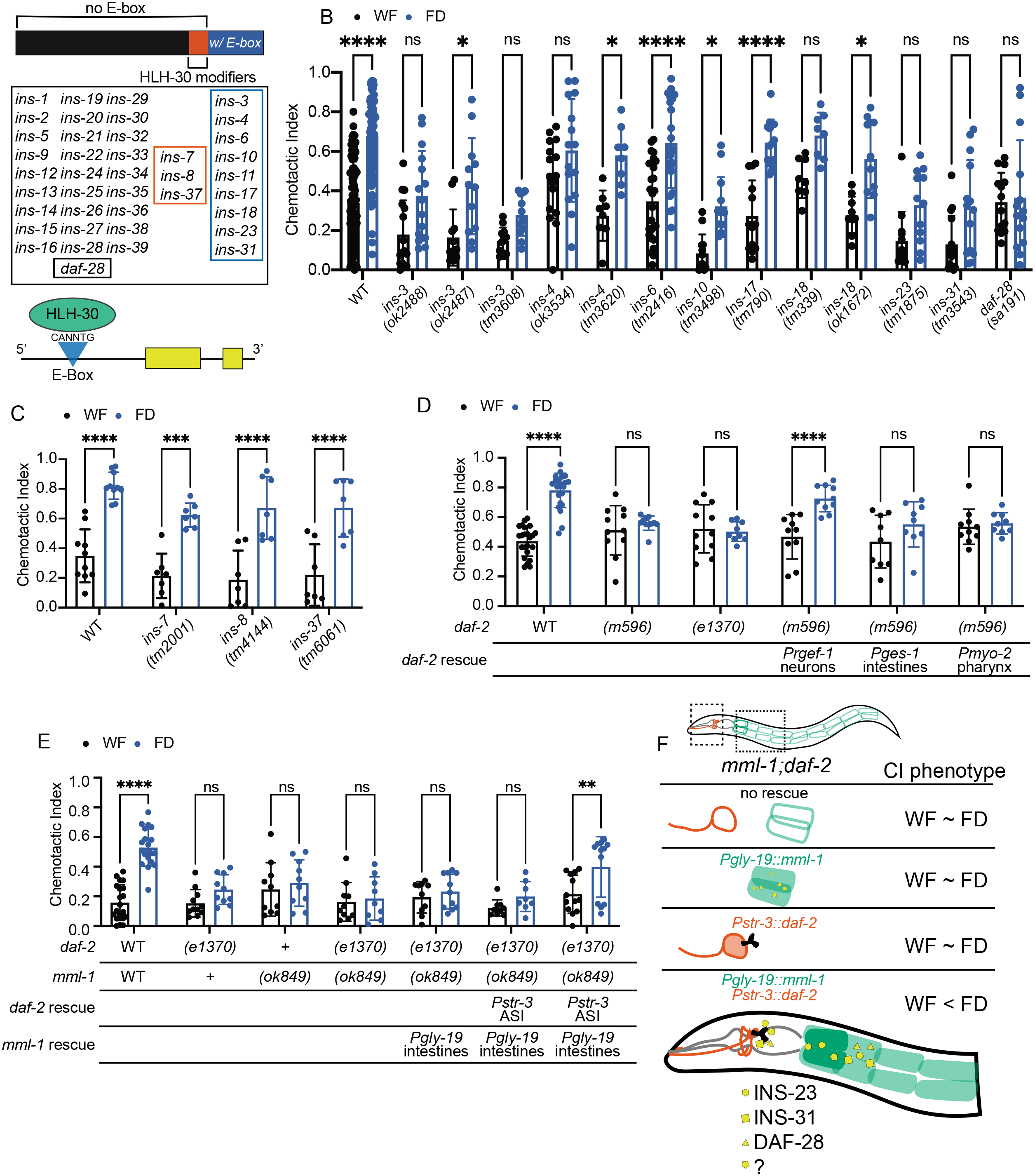
Sensory integration changes require HLH-30-regulated insulins and *daf-2* is required in neurons with *mml-1* in the intestine. (A) HLH-30 interacts with *C. elegans* insulin peptides. Of the 40 insulin-like peptides encoded in the *C. elegans* genome, 22% have an HLH-30 binding motif (CANNTG E-box, blue) in the 5’ UTR (< 300bp upstream of start site) [51]. 7% of insulins have been shown to regulate the localization of HLH-30 but do not contain an E-box (orange, “HLH-30 modifiers”). An illustration of a representative insulin peptide with two yellow exons and an upstream E-box with HLH-30 initiating transcription. (B) All insulins known to contain an HLH-30 binding motif in the 5’ UTR were tested using the standard sensory integration assay. When available, more than one allele was tested (N≥8) for each insulin, with wild-type (N2) animals tested with each mutant. (C) Insulins previously shown to regulate HLH-30 localization (*ins-7, ins-8, ins-37*) were tested using the standard sensory integration assay alongside wildtype (N2) control. N≥7. (D) *daf-2* mutants and tissue-specific rescues are tested in the standard sensory integration assay N ≥ 9 for each strain tested alongside wild-type N2. *daf-2* is rescued in neurons, intestines, and pharynx using tissue-specific promoters. (E) *daf-2, mml-1,* and *daf-2 mml-1* mutants were tested in standard sensory integration assays. *mml-1* was rescued in a tissue-specific manner in intestinal cells (*Pgly-19)* and *daf-2* was rescued in the ASI neurons (P*str-3*), alongside N2 controls. N≥10. (F) Schematic showing requirement of *mml-1* in the intestine, insulin-like peptides, and *daf-2* in ASI neurons. CI phenotype means Chemotactic Index phenotype, where wildtype animals displace a chemotactic index of WF < FD. All graphs were analyzed using a two-way ANOVA, determined to have significant differences across well-fed and food-deprived conditions. WF/FD comparisons were then performed as pairwise comparisons within each genotype or treatment as t-tests with Bonferroni’s multiple comparisons test. * p<0.5, ** p<0.01, *** p<0.001, **** p<0.0001, ns p>0.05.

Next, we probed the role of the insulin receptor, DAF-2, in affecting 3 hr-food deprivation evoked changes in sensory integration. Consistent with our analysis of mutants in various insulin-like peptide genes, we found that mutants in the insulin receptor, DAF-2, were also defective in their response to food deprivation (**Figure 5D**). To localize the site of DAF-2 action, we analyzed the effect of rescuing this receptor in different tissues. We found that expressing *daf-2* under neuronal, but not intestine or pharyngeal muscle promoters [57] restored normal behavior to the *daf-2* mutants (**Figure 5D**). Taken together, these results suggest that neuronally expressed DAF-2 receptors might detect INS-23, INS-31, and other insulin-like peptides released from the intestine, particularly those hindered by DAF-28 binding.

We then sought to test whether the upstream MML-1/Mondo A and the downstream DAF-2 insulin receptor function in the same pathway. We generated an *daf-2,mml-1* double mutant, which did not show any additional defects in the 3 hr food-deprivation evoked change in sensory integration when compared to either *mml-1* or *daf-2* single mutant (**Figure 5E**), suggesting that these two might function in the same pathway. We also found that expressing MML-1 in the intestine alone was not sufficient to restore wild-type behavior to the double mutant. In contrast, restoring MML-1 to the intestine and DAF-2 in ASI sensory neurons restored normal integration response after food deprivation (**Figure 5E**). Together, these data show that while MML-1 is required in the intestine, DAF-2 is required in ASI neurons and these genes act in the same pathway to alter food deprivation-modulated integration behavior (**Figure 5F**).

### ASI chemosensory neurons use insulin-signaling pathways to integrate intestine-released peptide signals

We then sought to identify components of the DAF-2 signaling pathway (**Figure 6A**) in ASI chemosensory neurons that were required to alter food-deprivation evoked change in sensory integration. We observed that mutants in the insulin-signaling pathway components including the FOXO family transcription factor *daf-16*, serine/threonine kinases AKT-1, AKT-2 (*akt-1*, *akt-2*), 3-phosphoinositide-dependent kinase 1 (*pdk-1*) and lipid phosphatase (*daf-18*, PTEN suppressor) performed normally in the sensory integration assay after food deprivation (**Figure 6B**) [58, 59]. In contrast, mutants in the phosphoinositide 3-kinase (PI3K, *age-1*) and Rictor (*rict-1)* [a key component of the mTORC2 complex [60, 61]] were defective in their copper sensitivity after food deprivation (**Figure 6B and 6C**). Similar to our *daf-2* rescue experiments, we found that Rictor was also required in ASI neurons to restore normal food-deprivation behavior to *rict-1* mutants (**Figure 6C**). These results suggest that PI-3 Kinase and Rictor might function in the same pathway downstream of DAF-2 receptors in ASI neurons. Previously, Rictor has been shown to act in the intestine to mediate an intestine-to-neuron signaling to affect dauer formation [62]. Our results show that Rictor can also act in ASI neurons as part of an intestine-to-neuron signal to alter sensory integration behavior. Collectively, we suggest that food deprivation engages DAF-2 signaling in ASI chemosensory neurons to alter the animal’s copper sensitivity allowing it to cross the copper barrier more readily.

**Figure 6:**
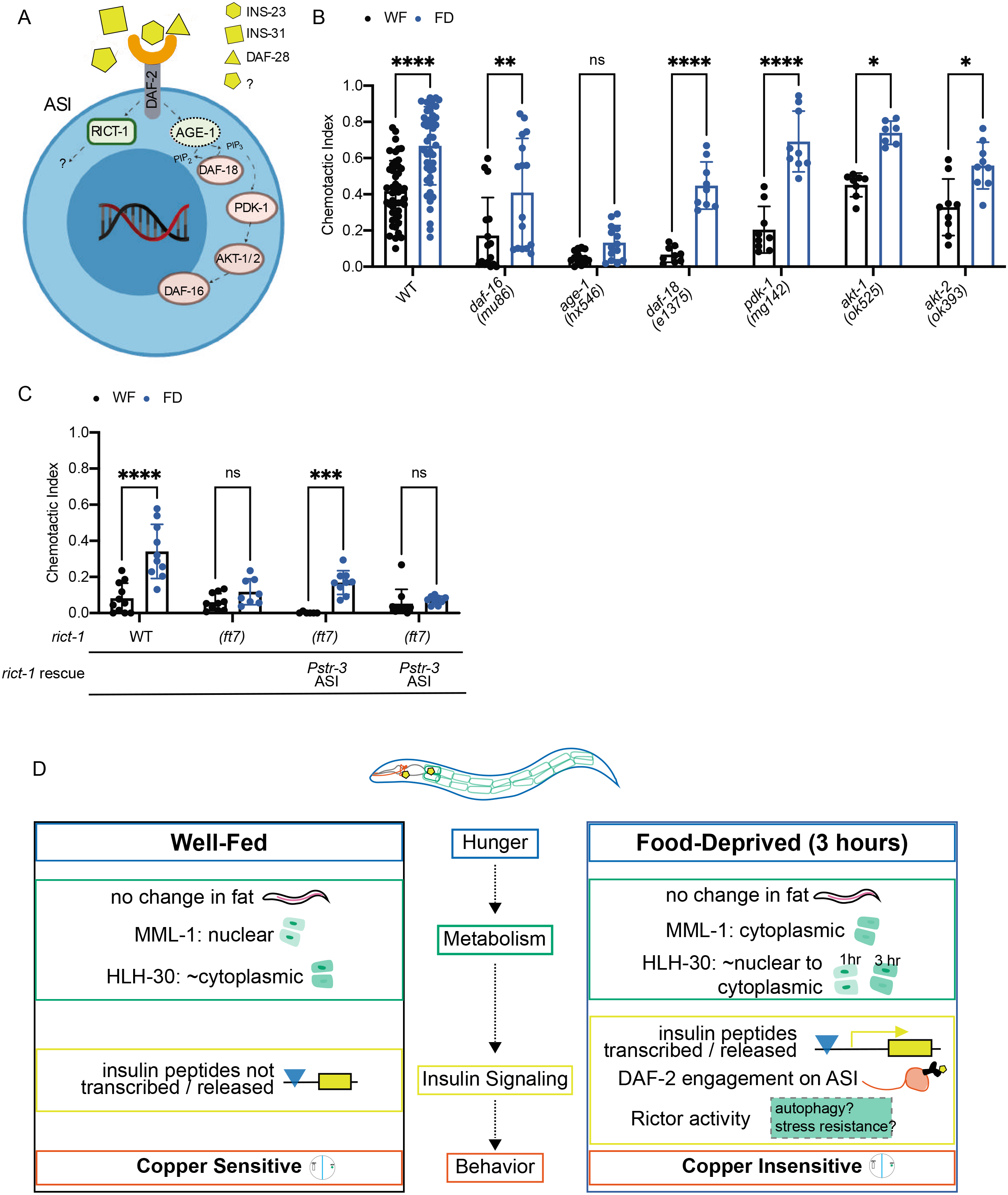
ASI chemosensory neurons use insulin-signaling pathway to integrate intestine-released peptide signals. (A) Schematic of an ASI neuron’s *daf-2*-mediated canonical and non-canonical insulin signaling. Summary of the findings in B and C. (B) Standard sensory integration assay performed with mutants in the canonical insulin signaling pathway (*daf-16, age-1, daf-18, pdk-1, akt-1,* and *akt-2*), alongside wild-type N2 N≥7. (C) Standard sensory integration assay performed with wildtype N2 (*rict-1* +), *rict-1* mutants, and *rict-1* mutants with *rict-1* rescued in ASI (*Pstr-3*). N ≥9. (D) Summary of data and proposed model through which food deprivation alters animal behavior. (B-C) were analyzed using two-way ANOVA, determined to have significant differences across well-fed and food-deprived conditions. WF/FD comparisons were then performed as pairwise comparisons within each genotype or treatment as t-tests with Bonferroni’s multiple comparisons test. * p<0.5, ** p<0.01, *** p<0.001, **** p<0.0001, ns p>0.05.

## Discussion

We used food deprivation in *C. elegans* as a model to understand how food deprivation modifies behavior. We show that food-deprived animals reversibly alter their behavior by reducing their repellent responsiveness, allowing them to traverse potentially toxic environments in their search for food. The *C. elegans* intestine sense the lack of food leading to cytosolic MML-1 and HLH-30, which in turn promotes the release of multiple insulins (INS-23, INS-31 and potentially others). These intestine-released peptides bind DAF-2 receptors and are processed by downstream PI-3 Kinase and RICT-1 in ASI and other neurons to reduce copper sensitivity and alter behavior (**Figure 6D**).

Multicellular animals’ sense and regulate glucose homeostasis at several levels. While insulin and glucagon maintain constant levels of circulating glucose, the Myc-family transcription factors are used within cells. Glucose uses cell membrane-localized transporters to enter cells, where it is rapidly converted into glucose-6-phosphate [63]. This intermediate metabolite is sensed by the Myc-Max complex, which binds glucose-6-phosphate and translocates to the nucleus where it regulates the transcription of glucose-responsive genes [41]. While the role of ChREBP/MondoA-Mlx-glucose-6-phosphate complex in regulating transcription is well studied [42, 64, 65], the role of these proteins in the cytoplasm remains poorly understood. We show a specific role for MML-1 (MondoA homolog), but not MXL-2 (Mlx homolog) in the intestine in reducing copper sensitivity after food deprivation. Additionally, we show that HLH-30, an ortholog of TFEB, is also required for attenuating copper sensitivity after food deprivation. Intriguingly, MML-1/MondoA and HLH-30/TFEB are both basic helix-loop-helix transcription factors and have been shown to act in concert to modify signaling networks and affecting global states like reproduction or survival [45]. Like MML-1/MondoA, a role for HLH-30/TFEB in the cytoplasm has also not been defined. We suggest that MML-1/MondoA and HLH-30/TFEB accumulation in the cytoplasm (in food deprived animals) enables the intestine to release peptide(s) relaying a “lack of glucose” signal to other tissues.

Helix-loop-helix transcription factors in *C. elegans* like MML-1/MondoA and HLH-30/TEFB are known to bind similar E-box elements (CACGTG) and have large overlap in their target gene expression [43, 45]. Additionally, previous studies have identified multiple insulin-like peptide genes whose expression is regulated by HLH-30 and other insulin-like peptide genes, which can affect the subcellular localization of HLH-30 [51, 54]. We screened this subset of insulin-like peptide genes to identify multiple candidates relaying food status signals from the intestine to other tissues. The *C. elegans* intestine has been previously shown to be a key tissue where the transcription of insulin peptide genes is regulated [50]. While we have not directly demonstrated that our candidate insulin-like peptides are released from the intestine, we speculate that food deprivation promotes their release relaying the “lack of food” signal.

We show that the ASI chemosensory neurons use the tyrosine kinase insulin receptor (DAF-2) to integrate these signals. Three lines of evidence suggest that the intestine is releasing multiple insulin-like peptide(s) - first, mutants in *ins-23*, *ins-31* and *daf-28* are defective in their food-deprivation evoked change in behavior, second, the insulin receptor (DAF-2) integrates these signals and third, MML-1 is required in the intestine and acts in the same pathway as the DAF-2 receptor, which acts in ASI neurons. We also define additional insulin signaling pathway components in ASI neurons. While FOXO (DAF-16), AKT kinase −1 and −2, PDK-1 and PTEN (DAF-18) are not required, we show that AGE-1 (PI-3 Kinase) and Rictor (a component of the mTORC2 complex) are required to integrate intestine-released peptide signals. While PI-3 kinase has been shown to act via AKT kinase to activate the Rictor [60, 61], our result hint at an AKT kinase-independent mechanism for PI-3 Kinase to signal to Rictor and the mTORC2 complex.

Multiple studies have also highlighted the role of insulin signaling in relaying starvation-related signals to various neurons. Starvation has been shown to decrease the secretion of INS-18 from the intestine, which antagonizes DAF-2 receptor in ADL neurons and modifies pheromone-mediated behaviors [20]. Also, starvation has been shown to be associated with increased octopamine signaling, which transforms CO_2_ attraction to repulsion in starved animals [18]. Moreover, starvation has also been shown to recruit ASG neurons to cooperate with ASE neurons and drive avoidance to high salt [19]. We speculate that our intestine-to-neuron insulin signaling pathway leads to altered ASI function and altered copper sensitivity. Consistently, starvation has been shown to increase ASI neural activity in response to food-stimuli [66]. These data are also consistent with previous studies showing that ASI neurons playing a crucial role in modifying behavior after 6 hours of food deprivation [17, 67]. Taken together, we speculate that food deprivation leads to an increase in insulin signaling from the intestine to ASI neurons, which alters neuronal activity and reduces the animal’s sensitivity to copper, allowing it to cross the barrier more readily. More broadly, these studies link transcription factors and insulin signaling from the intestine to neurons to modify sensory behavior, a mechanism likely conserved across species.

## Methods

### Strains

*C. elegans* strains were grown and maintained under standard conditions [68]. All strains used are listed in **Supplementary Table 1**.

### Behavior Assays

All animals were grown to adulthood on regular nematode growth medium (NGM) plates seeded with OP50 (OD_600_ ∼ 0.2) before they were washed and transferred to new food (standard NGM plates seeded with OP50) or food-free plates (standard NGM plates) respectively for the indicated duration. Sephadex beads (G-200) were added to both the empty NGM plate and the OP50 lawn in experiments for Figure 3E-F. Sensory integration assays were performed on 2% agar plates containing 5 mM potassium phosphate (pH 6), 1 mM CaCl_2_ and 1 mM MgSO_4_, made the day before the experiment. Repellent gradients (including CuSO_4_ (Copper (II) sulfate pentahydrate, Sigma 209198), glycerol (, NaCl (, fructose (D-(-) Fructose Sigma F0127), and quinine (Sigma 22620)) were established by dripping 25 μl of solution, dissolved in water, across the midline of the plate [30]. This solution was allowed to dry overnight. Prior to the assay, the animals were washed from the food or food-free plates into Eppendorf tubes. Each treatment group was serially washed once with M9+MgSO_4_ and 3 times with Chemotaxis buffer (5 mM potassium phosphate (pH 6, Fisher BP362 monobasic and Fisher BP363, dibasic), 1 mM CaCl_2_ (Sigma C1016) and 1 mM MgSO_4_ (Sigma M7506)) before being transferred to the assay plates. Glass Pasteur pipets were used to prevent loss of animals due to sticking in plastic pipette tips. Immediately after plating 100-200 animals in a small drop of chemotaxis buffer, 1 μL of attractant with 1 μL of 1M sodium azide in water (Sigma 71289) was placed on the opposite side of the chemotaxis plate. Attractants used were diacetyl (2,3-Butanedione Sigma 11038), Isoamyl alcohol (3-methyl-1-butanol, Sigma 77664), and Benzaldehyde (Sigma 418099) diluted in Ethanol. If necessary, the small drop of animals was dabbed gently with the edge of a Kim wipe and the lid was immediately replaced. After 45 minutes or at indicated times, the integration index was computed as the number of worms in the odor half of the plate divided by the total number of animals on the plate. For each experiment, at least two plates were tested each day with experiments performed on at least three different days. Unless otherwise noted, the repellant is a dried stripe of 25 μL 50 mM CuSO_4_ (Copper (II) sulfate pentahydrate, Sigma 209198) in water and the attractant is 1 μL 0.2% diacetyl (2,3-Butanedione Sigma 11038) diluted 1:500 in 100% ethanol.

### Statistics

For sensory integration, experiments were performed at least 3 times with at least 2 plates per genotype/condition (unless otherwise noted). For strains with extrachromosomal arrays, only animals expressing the co-injection markers were counted. Every condition was performed with N2 (wild-type) controls at the same time. Two-way ANOVAs with post-hoc Bonferroni-corrected multiple comparisons were performed across WF/FD conditions, only if the factor was significant. For all figures, p values are represented by: * p<0.05, ** p<0.01, *** p<0.001, **** p<0.0001.

### Single animal avoidance assay: Copper drop test

Experiments were performed as previously described [31]. Animals are moved from a food to a food-free assay plate. A capillary tube is used to deliver a drop of test compound (1.5 mM CuSo_4_) 0.5 - 1 mm away from the head of the animal and its responses scored. Positive avoidance indicates an animal executing a large reversal and omega bend within 3 seconds of sensing the test compound. Five animals are tested per condition and with each animal exposed to 10 drops and the percent avoidance is plotted. Assay is replicated at least three times by an investigator who is blind to the conditions being tested.

### Tracking

Sensory integration behavior assays using 50 mM CuSO_4_ (Copper (II) sulfate pentahydrate, Sigma 209198) in water and 1 μL 0.2% diacetyl (2,3-Butanedione Sigma 11038) attractant (1:500 in 100% Ethanol) were performed with well-fed and food-deprived animals. Animal behavior was recorded for 45 minutes using a Pixelink camera (1024×1024 pixels at 3 frames per second). The imaging field of view was approximately 47 mm x 47 mm. WormLab software (MBF Bioscience) was used to identify and track the midpoints of worms in each video. Custom MATLAB software (https://github.com/shreklab/Matty-et-al-2021) was used to further clean the data (i.e. remove putative tracks that did not correspond to animal behavior) and analyze individual tracks. Tracks were excluded if they met any of the following criteria: 1) overlapped with shadows or markings; 2) lasted less than 10 seconds; 3) travelled fewer than 30 pixels^2^; or 4) traveled less than 10 pixels in any direction. Valid animal tracks were then plotted (**Figure 2A, 2B**) and analyzed as described below.

### Tracking analysis

The number of animals in each experiment is estimated from the maximum number of simultaneous tracks identified in a single frame. An average of 33.2 ± 10.4 animals were assayed across all conditions. Because the field of view does not encompass the entire plate, the number of tracks identified in each frame decreases over time as animals crawl to other regions of the plate. To quantify the number of animals crossing the copper barrier as a function of time, the number of unique tracks that started past the copper barrier was divided by the number of unique tracks in the entire field-of-view. This fraction of cumulative unique tracks that crossed the copper barrier was calculated for 15, 30, and 45 minutes (**Figure 2C, 2F, 2I; Supplementary Figure 3A**). To better understand animals’ avoidance of copper and attraction to diacetyl, the probability of an animal residing at a particular distance from the barrier was calculated for 1 mm bins. The total number of tracked midpoints at each time point located in each 1 mm bin was summed and divided by the total number of tracked midpoints across all bins (**Figure 2D, 2G, 2J; Supplementary Figure 3C**). Additionally, animal velocity was calculated by computing the Euclidean distance of a track over a 2 second window. In each video, the average velocity of all tracks was computed as a function of distance from the copper barrier in 1 mm bins (**Figure 2E, 2H, 2K; Supplementary Figure 3B**).

### Visualizing Copper Gradients

Copper sulfate gradients were visualized using 1-(2-Pyridylazo)-2-naphthol (PAN, Sigma 101036). Plates with 25 μL of 5 mM, 25 mM, 50 mM and 100 mM CuSO_4_ dripped down the midline were dried overnight. 1 mL of 0.01% PAN indicator was added to plates the next day and allowed to dry. The plates with PAN indicator were incubated overnight and imaged the following day to allow for saturation of the signal. Images and quantification of the copper barrier is shown in **Supplementary Figure 1**.

### Fat quantification

Oil red O staining was conducted as previously described [36]. Briefly, 10-20 N2 adults were allowed to lay eggs for 1 hour on NGM plates seeded with OP50. The adults were removed and eggs were allowed to develop for 3 days. These day-1 adult animals were either removed from food and placed on an empty NGM plate for 3 hours or 6 hours or placed on a new plate with OP50 food. 5 mg/mL Oil Red O (Sigma, O9755) in 100% isopropanol was prepared as a working solution and diluted 3:2 in 60% isopropanol on the day before use. Mixture was kept from the light and filtered using a 0.2 μm cellulose acetate syringe filter and allowed to mix on a rocker overnight. Animals were washed off plates with PBST (PBS + 0.01% Triton X-100 (Sigma, X100)at the appropriate times and washed once. Animals were fixed in 40% isopropanol and shaken at room temperature for 3 minutes. Isopropanol was removed and 600 μL of the Oil Red O diluted solution was added to each tube. Each tube was nutated for 2 hours at room temperature, away from light. Animals were washed once with PBST and nutated for another 30 minutes. Animals were washed once more and prepared for imaging. Approximately 20 worms from each treatment group were pipetted onto a microscope slide and covered with a coverslip. Images were collected on upright Zeiss Axio Imager.M2 at 10X using an AxioCam 506 Color camera. Images were quantified using color deconvolution in ImageJ, normalized to background and an unstained region of an animal. Within each experiment, the same thresholds were used across treatments. Approximately 20 animals were quantified within each condition on each experimental day, performed across three different days.

### Imaging

Transgenic animals (HLH-30::GFP and MML-1::GFP) were grown to day 1 adulthood (3 days post hatching) via a one-hour hatch off on standard NGM plates seeded with OP50. Animals were picked onto empty NGM plates for 1, 2, and 3 hours for food deprivation or placed on a new NGM plates with OP50. Animals were picked onto thin agar pads on microscope slides and anesthetized with 100 μM tetramisole hydrochloride (Sigma-Aldrich L9756) immediately prior to imaging. Animals were imaged at 10X using an upright Zeiss Axio Imager .M2. At least 12 animals per group on three different days were imaged and qualitatively analyzed for localization to primarily cytoplasmic, nuclear, or both in intestinal cells, with the investigator blind to food deprivation status.

### Molecular Biology and Transgenics

The following primers were used for amplifying full-length cDNAs:

*mml-1*

forward 5 ‘TATTTAGCTAGCATGTCGCGCGGGCAGATTATACACAG reverse 5’CGGGGTACCGAGCAGTTCAAAATGGATTTTTGAGTTGTTGC

*rict-1*

forward 5’TATTTAGCTAGCATGGACACTCGTCGAAAAGTGTATCAC reverse 5’CGGGGTACCTAAAAGATTTGCTGCAGGAATGCTCTCG

*daf-2*

forward 5’ TATTTAGCTAGCAATGAATATTGTCAGATGTCGGAGACGA 3’ reverse 5’ CGGGGTACCTCAGACAAGTGGATGATGCTCATTATC 3’

cDNAs corresponding to the entire coding sequences of *mml-1, rict-1,* and *daf-2* genomic region were amplified by PCR using primers above and expressed under tissue or cell selective promoters.

Tissue specific expression was achieved with *Prgef-1* for neurons, *Pges-1* and *Pgly-19* for the intestine, and *Pmyo-2* for pharynx, [69–74]. Cell-specific expression used *Pstr-3* was used for cell-specific expression in ASI neurons [75]. For all experiments, a splice leader (SL2) fused to *mCherry* or *gfp* transgene was used to confirm expression of the gene of interest in either specific cells or tissues. Germline transformations were performed by microinjection of plasmids [76] at concentrations between 50 and 100 ng/μl with 10 ng/μl of *elt-2::gfp* as a co-injection marker.

## Author Contributions

M.A.M. and H.E.L. conceived and conducted the experiments, interpreted the data, and co-wrote the paper. A.S., A.C., and K.K., conducted behavioral assays; A.S.’ experiments were conducted in M.H.’s laboratory. J.A.H. analyzed tracking data. S.H.C. conceived the experiments, interpreted the data and co-wrote the paper. All co-authors provided feedback on the manuscript.

## Acknowledgements

We thank A. Dillin, T. Ishihara, S. Lockery, A. Samuelson, E. Troemel, M. Zhen, the National BioResource Project (NBRP, Japan) and Caenorhabditis Genetics Center (CGC) for strains; C. Bargmann, E. Hallem, M. Hilliard, A. van der Linden, P. McGrath, D. Pilgrim and P. Sengupta for constructs; S. Srinivasan and lab members for RNAi clones and help with Oil-Red O staining; and M. Tamés, Z. Liu, and C. Yang for technical help with behavioral and imaging studies. We are also grateful to Jing Wang, and members of the Chalasani lab for critical comments, advice, and insights. This work was funded by grants from The Rita Allen Foundation, The W.M. Keck Foundation and NIH R01MH096881 to S.H.C., NSF Postdoctoral Research Fellowships in Biology Program Under Grant No. 2011023 (M.A.M), Socrates Program (H.E.L. Award #NSF-742551) and Graduate Research Fellowship from NSF (H.E.L., and J.A.H.).

## Supplementary Material Legends

**Supplementary Figure 1 (to accompany Figure 1):**
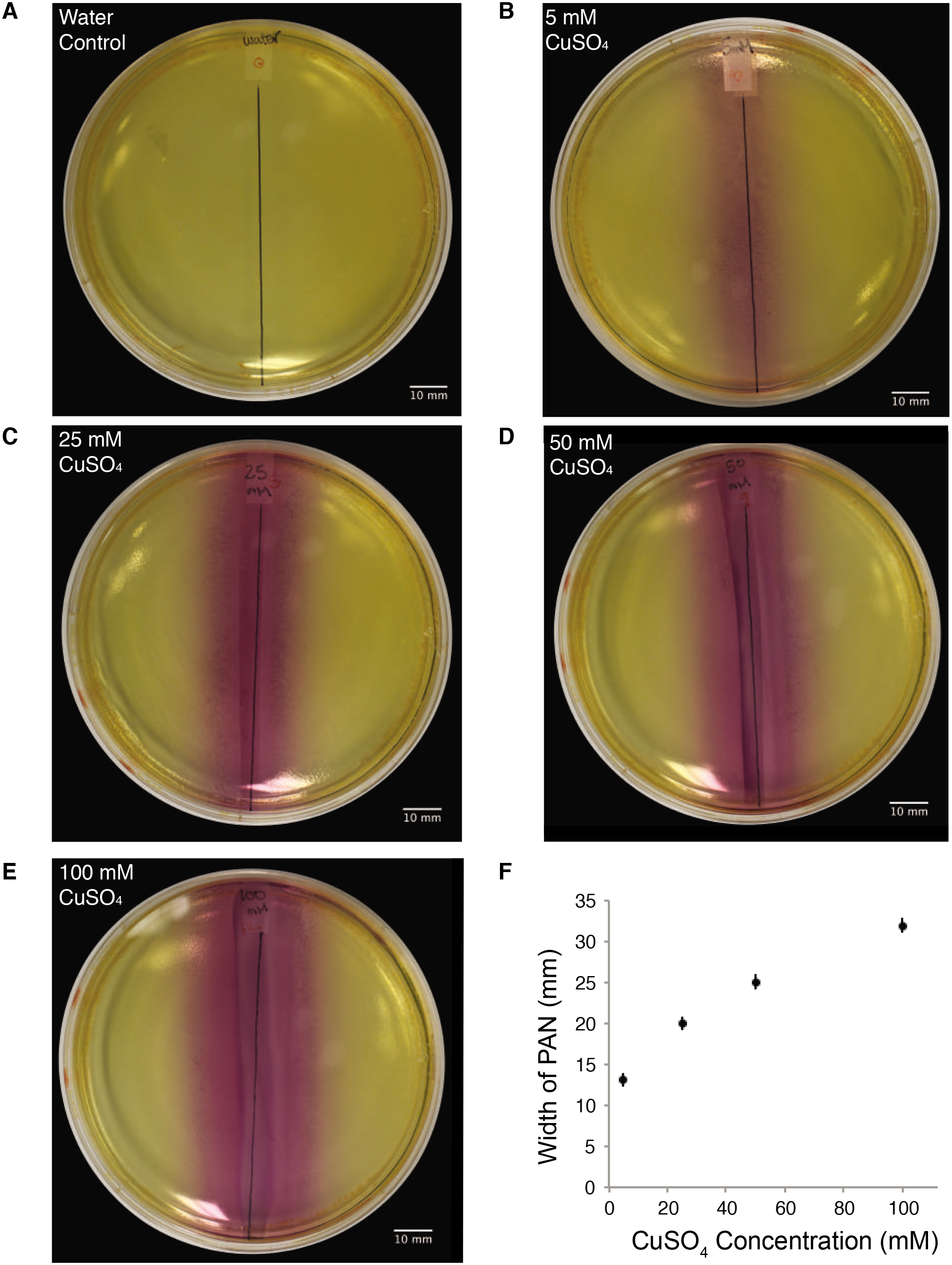
Spread of copper sulfate CuSO_4_ on agar plates visualized using 1-(2-pyridylazo)-2-naphthol. (A-E) 25 μl of (A) water as control, (B) 5 mM CuSO_4_, (C) 25 mM CuSO_4_, (D) 50 mM CuSO_4_, and (E) 100 mM CuSO4 was dripped and dried overnight along the midline of the plate to form a copper gradient. PAN indicator (1-(2-pyridylazo)-2-naphthol) distributed over the entire plate shows a gradient of orange-red upon chelation with copper ions. (F) Measured width of colored area with each data point representing the average width, error bars indicate SEM. N = 9.

**Supplementary Figure 2 (to accompany Figure 1):**
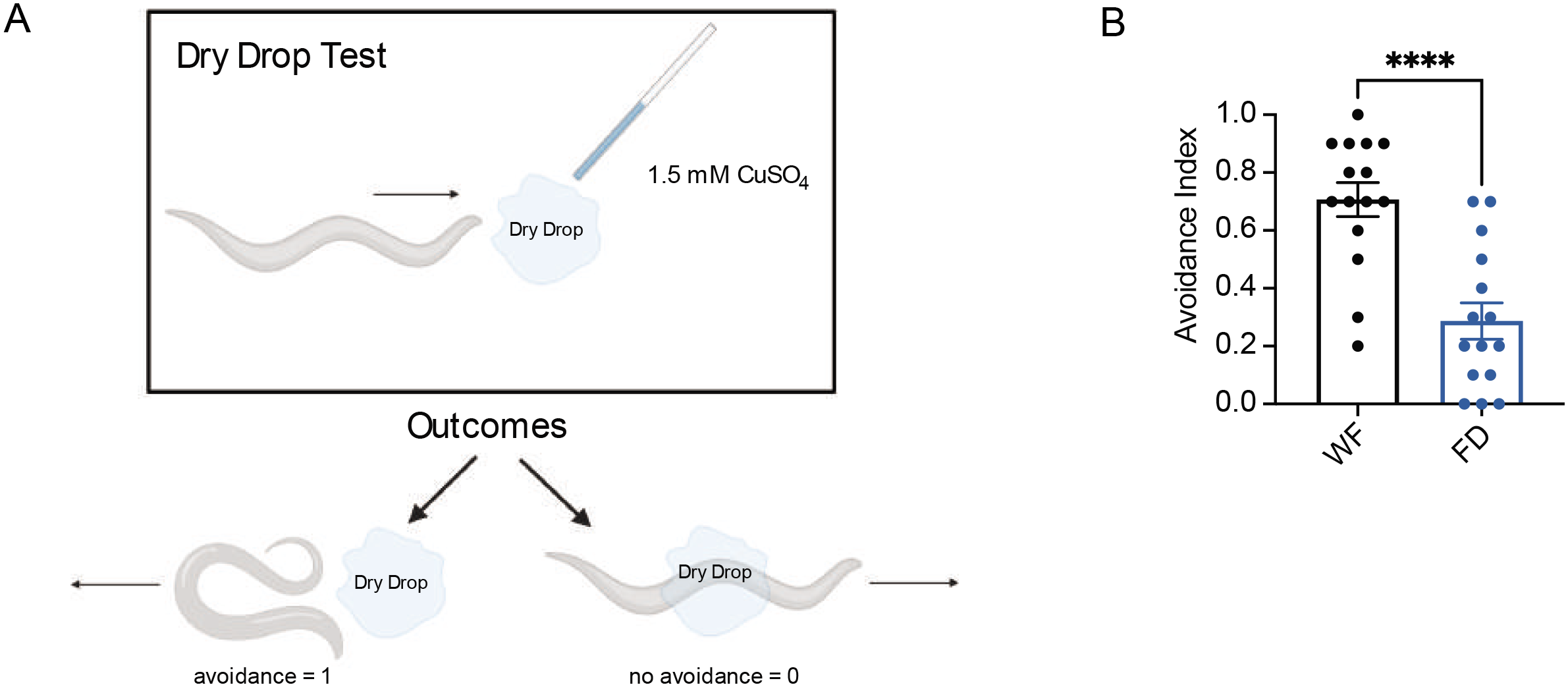
Food-deprived animals fail to avoid copper in single animal drop test. (A) Schematic for dry drop test shown in B. ∼300 nL of 1.5 mM CuSO4 is dropped ∼1 mm away from the animal’s forward motion. Turning away or backing up is considered “avoidance” and given a score of 1. Heading toward the dried drop is considered “no avoidance” and given a score of 0. (B) Quantification of the dry drop test. Food-deprived (FD) animals were starved for 3 hours. Each dot represents the average of ten trials (drops) for a single animal, N=15. Analyzed with an unpaired t-test * p<0.5, ** p<0.01, *** p<0.001, **** p<0.0001, ns p>0.05. Error bars are S.D.

**Supplementary Figure 3 (to accompany Figure 2):**
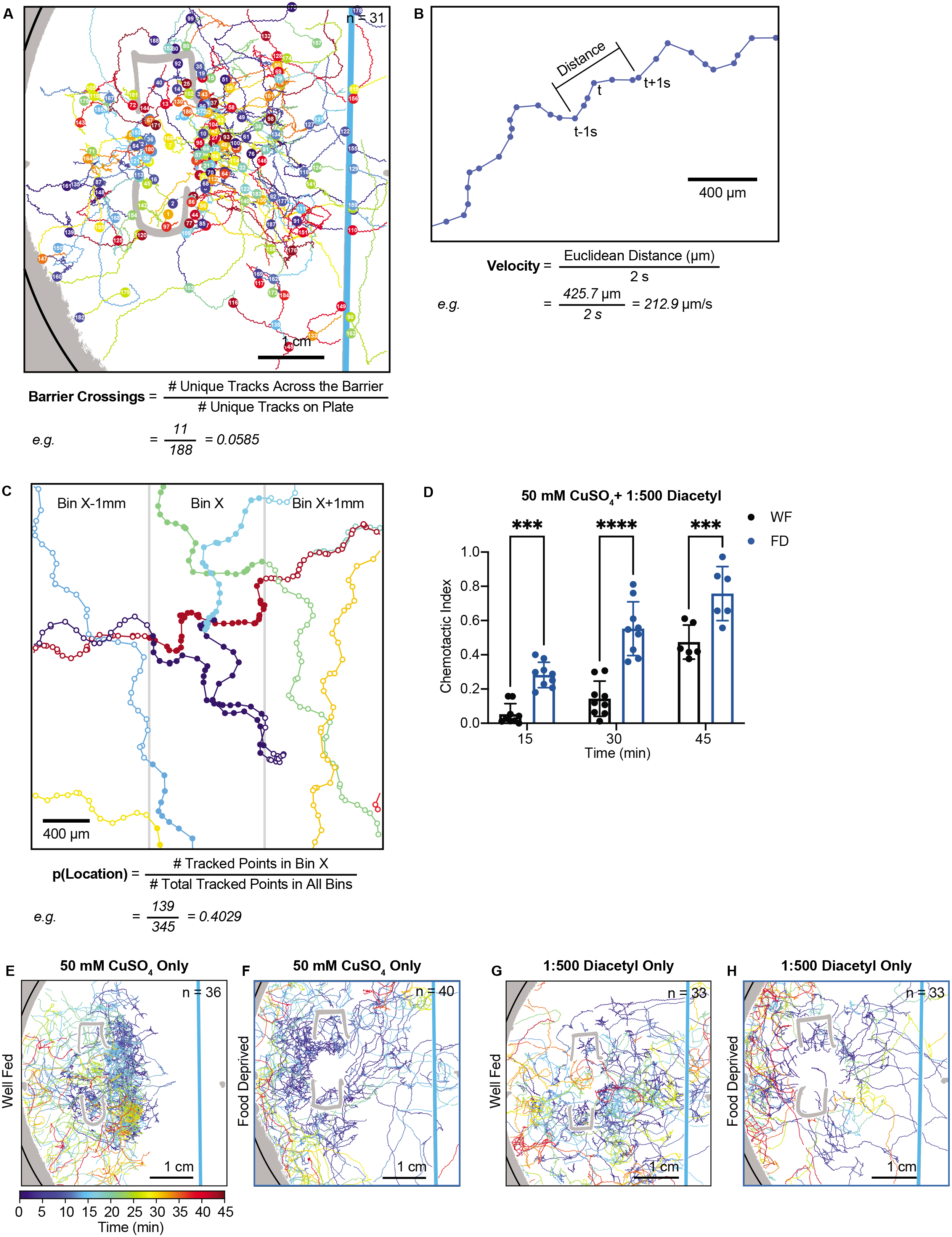
Description of measurements to define tracking dynamics and additional treatment groups. (A) Measuring Barrier Crossings. Worm tracks (n = 31) are plotted for the first 15 minutes of a representative sensory integration assay. 188 tracks are plotted in a unique color with the start of each track labelled with a numbered, circular marker. The number of unique, continuous tracks that started past the copper barrier was divided by the number of unique tracks in the entire field-of-view to obtain a measure of barrier crossing. In the example experiment shown, 11 unique tracks crossed the barrier out of 188 total tracks, resulting in a Barrier Crossings score of 0.0585 for this experiment after 15 minutes of recording. (B) Measuring Velocity. 10 seconds (i.e. 30 frames) of a single example worm track is plotted. The midpoint positions of the worm at each frame as identified by WormLab are plotted as filled circles connected by lines. For each time *t*, the velocity was calculated by computing the Euclidean distance of the track from time *t – 1* second to time *t + 1* second and dividing by the length of time, 2 seconds. Because these videos lack the special resolution necessary to accurately estimate absolute path length (and thus body bends), Euclidean distance is used. In the example given, the Euclidean distance of the 2 second time window centered at time *t* was 425.7 μm resulting in an instantaneous velocity of 212.9 μm/s. Velocity was calculated for every time point in this way. (C) Measuring Probability of Location. 9 unique worms tracks are plotted in a 3 mm x 3 mm field-of-view, a 9 mm^2^ inset of a 45-minute example experiment. The midpoint positions of the worms at each frame are plotted as circles connected by lines. Midpoints located in Bin X (1 mm wide) are represented by filled circles while midpoints located in the neighboring bins (Bin X-1 and Bin X+1, each 1 mm wide) are represented by open circles. The probability of a worm being located in Bin X is calculated by dividing the number of tracked midpoints in Bin X by the total number of tracked points in all bins. In the small example area shown, there are 139 points in Bin X and a total of 345 points across all 3 bins resulting in a p(Location) score of 0.4029. In the entire field of view there are 45 bins, yielding an average p(Location) score of 0.0222. This analysis was used in Figure 2D, 2G, 2J. (D) Graph of the chemotactic index (# animals on odor side / total # of animals) over time (15, 30, 45 minute bins). Well-fed (WF) animals appear with black dots and food-deprived (FD) animals are indicated with blue dots. Each dot represents a single plate of animals, with each plate measured at each time point (matched). Analyzed using a Two-Way ANOVA, determined to have significant differences across well-fed and food-deprived conditions. WF/FD comparisons were then performed as pairwise comparisons within each time period as t-tests with Bonferroni’s correction for multiple comparisons. * p<0.5, ** p<0.01, *** p<0.001, **** p<0.0001, ns p>0.05. (E) Worm tracks (n = 36) are plotted for a representative sensory integration assay of well-fed worms behaving in the presence of 50 mM CuSO_4_ in water (blue stripe) with no attractant (location not shown). Regions of the plate that were not able to be tracked are in gray with the edge of the plate indicated in black. Tracks are plotted and color coded for time. (F) Worm tracks (n = 40) are plotted for a representative sensory integration assay of 3 hour food-deprived worms. Conditions and plotting the same as in E. (G) Worm tracks (n = 33) are plotted for a representative sensory integration assay of well-fed worms behaving in the presence of no barrier (blue stripe) with attractant is 1 μL 0.2% diacetyl (1:500) in 100% ethanol (location not shown). Regions of the plate that were not able to be tracked are in gray with the edge of the plate indicated in black. Tracks are plotted and color coded for time. (H) Worm tracks (n = 31) are plotted for a representative sensory integration assay of 3 hour food-deprived worms. Conditions and plotting the same as in G.

**Supplementary Table 1: All worm strains used in the experiments.** Strain ID, genotype/allele, and how it is referenced in the paper is provided. If the strain is first described here (all IV strains), the method of creation is provided.

**Supplementary Table 2: All p-values for** figures 2D**, 2G, 2J and 2E, 2H, 2K.** The p-values shown are the result of multiple unpaired t-tests with Welch’s correction with Holm-Šídák post-hoc tests correction for multiple comparisons. Adjusted p-values are shown, with yellow shading for adjusted p-values <0.05, same shading as in figures 2D, 2G, 2J and 2E, 2H, 2K.

**Supplemental Movie S1. Sensory integration behavior of well-fed animals.** ∼150 Well-fed wild-type animals are placed in the standard sensory integration assay. Bracket indicates origin where animals are placed, spot shows position of 1:500 diacetyl odor, midline indicates repellent CuSO_4_ barrier.

**Supplemental Movie S2. Sensory integration behavior of food-deprived animals.** ∼150 Wild-type animals’ food-deprived for three hours are placed in sensory integration behavior assay. Bracket indicates origin where animals are placed, spot shows position of 1:500 diacetyl odor, midline indicates CuSO_4_ barrier.

